# Periosteum-derived podoplanin-expressing stromal cells regulate nascent vascularization during epiphyseal marrow development

**DOI:** 10.1101/2021.07.05.451096

**Authors:** Shogo Tamura, Masato Mukaide, Yumi Katsuragi, Wataru Fujii, Koya Odaira, Nobuaki Suzuki, Shuichi Okamoto, Atsuo Suzuki, Takeshi Kanematsu, Akira Katsumi, Akira Takagi, Katsue Suzuki-Inoue, Tadashi Matsushita, Tetsuhito Kojima, Fumihiko Hayakawa

## Abstract

Bone marrow development and endochondral bone formation occur simultaneously. During endochondral ossification, periosteal vasculatures and stromal progenitors invade the primary avascular cartilaginous anlage; this induces primitive marrow development. We previously determined that bone marrow podoplanin (PDPN)-expressing stromal cells exist in a perivascular microenvironment, and promote megakaryopoiesis and erythropoiesis. In this study, we aimed to examine the involvement of PDPN-expressing stromal cells in the postnatal bone marrow generation. We found that periosteum-derived PDPN-expressing stromal cells regulate vascularization during postnatal epiphyseal marrow development. Our findings suggest that these cells act as pericytes on the primitive vasculature of the nascent marrow. They invade the cartilaginous epiphysis and regulate marrow development and homeostasis by maintaining vascular integrity. To the best of our knowledge, this is the first study to comprehensively examine how PDPN-expressing stromal cells contribute to marrow development and homeostasis.

## Main

The bone marrow is a three-dimensional tissue within the bone cavity that is composed of vasculature, extracellular matrices (ECMs), and stromal cells^1, 2^. Bone marrow development and bone formation occur simultaneously. In mammals, bones are formed via two distinct mechanisms, i.e., intramembranous and endochondral ossification. Intramembranous ossification is the process of bone development from soft connective tissue that is involved in the formation of flat bones of the skull (calvarial bones and mandibles) and part of the clavicles. Endochondral ossification is the process of bone development from cartilage that forms all bones of the body, except the flat bones of the skull. During the endochondral ossification, the vascular invasion of the primary avascular cartilaginous anlage triggers the formation of an embryonal primary ossification center (POC) and postnatal secondary ossification center (SOC) in the diaphysis and epiphysis, respectively^3^. The invading vasculature transports chondroclasts, osteoblast progenitors, and stromal progenitors from the periosteum to the POC or SOC^4, 5^. The invading vasculature and stromal progenitors generate the primitive marrow inside the bone cavity.

Skeletal stem cells (SSCs) are a heterogenous population of stromal cells that play a role in bone marrow generation, skeletal tissue development, homeostasis, and regeneration^6, 7^. In mice, SSCs exist within the periosteum (P-SSCs)^8–11^, bone marrow (BM-SSCs, also known as bone marrow stem/stromal cells, BMSCs)^12–23^, and growth plate resting zone (GP-SSCs)^24, 25^. Studies involving mouse BM-SSCs have identified several BM-SSC subpopulations [e.g., CXCL12 abundant reticular cells (CAR cells)^12–15^, leptin receptor-positive cells^16–18^, *nestin*^GFP^-positive cells^21, 22^, *grem1*^Cre-ERT^-positive cells^20^, and Mx1-positive cells^11, 19^]. These BM-SSC subsets are present at the abluminal surface of blood vessels and induce the formation of hematopoietic microenvironments via the expression of hematopoietic regulators, such as CXCL12 and SCF^26, 27^. Previously, we have identified podoplanin (PDPN, also known as gp38 or T1a)-expressing stromal cells that existed in the bone marrow^28^. In adult mice, PDPN-expressing stromal cells induce the generation of a perivascular microenvironment that promotes megakaryopoiesis and erythropoiesis^28, 29^. However, the cellular sources and physiological functions of PDPN-expressing stromal cells during postnatal bone marrow development have not been elucidated.

The aim of this study was to characterize the cellular features of marrow PDPN-expressing stromal cells and disclose how these cells are involved in the postnatal bone marrow generation. To the best of our knowledge, this is the first study to investigate the role of PDPN-expressing stromal cells on marrow development and homeostasis. These study findings will improve our understanding of how stromal cells regulate the nascent bone marrow development and homeostasis.

## Results

### PDPN-expressing periosteal cells invade into the postnatal primary epiphysis and are present in primitive vascular beds

We and other research group (Baccin C et al) had previously detected PDPN-expressing stromal cells in the diaphyseal marrow of adult mice (> 8-week old)^28, 30^. In the present study, to investigate the source of PDPN-expressing stromal cells and their physiological functions in the postnatal nascent bone marrow, we attempted to screen their distribution in mice femurs at postnatal day 21 (P21). Because the number of PDPN-expressing stromal cells in the marrow was very low, we enriched PDPN-positive marrow cells using magnetic microbeads (Fig. 1a). Using flow cytometric analysis, we detected the PDPN-expressing stromal cells in the diaphyseal and epiphyseal marrow (Fig. 1b). In the stromal population of the PDPN-positive enriched marrow [Lin(-)CD31(-)CD45(-)CD51(+)CD150(-)], more PDPN-expressing stromal cells were detected in the epiphysis than in the diaphysis. The diaphyseal PDPN-expressing stromal cells were stem cell antigen-1 (Sca-1)-negative; these findings were consistent with those of our previous study^28^. In contrast, the epiphyseal PDPN-expressing stromal cells were mainly Sca-1 positive. These data indicate the characteristic differences between the epiphyseal and diaphyseal PDPN-expressing stromal cells. To analyze these newly identified cells, we evaluated the epiphyseal marrow using histological analysis. At P21—in the cryosection of the epiphysis—PDPN-expressing stromal cells were detected in the SOC and surrounding primitive vasculatures (Fig. 1c). Upon evaluating the expression of pericyte markers, platelet-derived growth factor receptor β (PDGFRβ), neuron-glial antigen 2 (NG2), and alpha smooth muscle actin (αSMA)^31^, in these cells, we found that the epiphyseal marrow PDPN-expressing stromal cells were PDGFRβ positive, NG2 positive, and αSMA negative (Fig. 1d-f). The pericytes, determined to be PDGFRβ(+)NG2(+)αSMA(-), are generally found on the capillary bed^32^. These findings indicate that epiphyseal PDPN-expressing stromal cells can act as pericytes in primitive capillary-like vascular beds in the developing SOC.

**Figure. 1.**
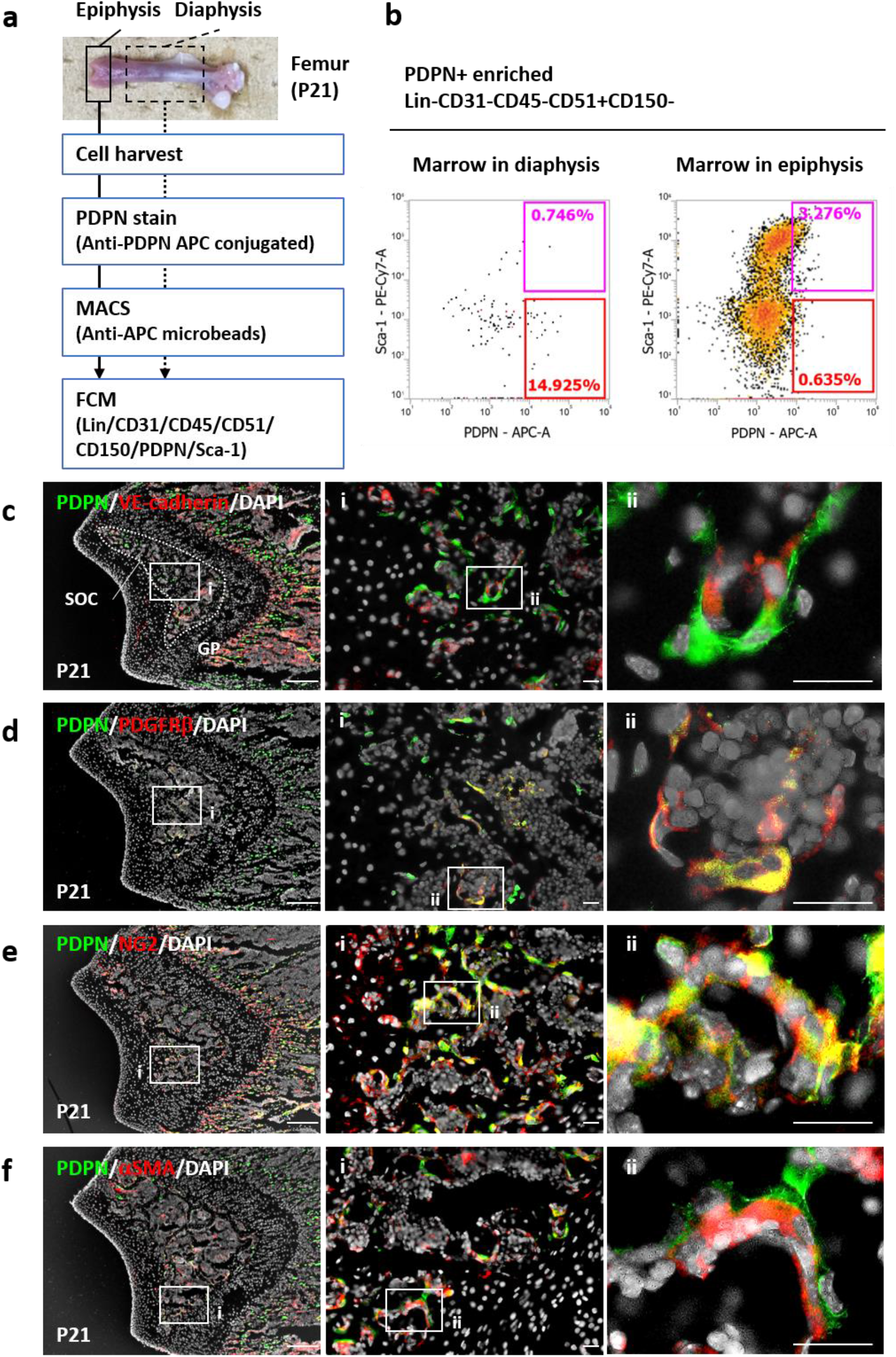
Epiphyseal PDPN-expressing stromal cells are the pericytes on the primitive capillary-like vascular bed of the secondary ossification center. (A) Scheme to detect marrow PDPN-expressing stromal cells via flow cytometry. (B) Representative flow cytometric data of PDPN-expressing stromal marrow cells in the epiphysis and diaphysis in mice at P21. PDPN-positive enriched marrow cells were analyzed by flowcytometry. Magenta and red gates indicate the PDPN(+)/Sca-1(+) and PDPN(+)/Sca-1(-) populations, respectively. The percentages inside each gate indicate the cells contained in the population of Lin(-)CD31(-)CD45(-)CD51(+)CD150(-). (C) Representative IHC images of P21 mouse epiphysis. Epiphysis cryo-sections were stained with PDPN, VE-cadherin, and DAPI. PDPN-expressing stromal cells surrounded primitive vasculatures. (D-F) Representative IHC images of P21 mouse epiphysis with PDPN/PDGFRβ/DAPI (D), PDPN/NG2/DAPI (E), and PDPN/αSMA/DAPI (F). Scale bars in the lefts indicate 200 μm. Scale bars in the middle and right panels indicate 50 μm. PDPN: podoplanin, P21: postnatal day 21, IHC: immunohistochemistry, VE-cadherin: vascular endothelial-cadherin, DAPI: 4′,6-diamidino-2-phenylindole, PDGFRβ: platelet-derived growth factor receptor-β, NG2: neuron-glial antigen-2, αSMA: α-smooth muscle actin

Next, we investigated how PDPN-expressing stromal cells populated the epiphyseal marrow. We sequentially chased epiphyseal SOC development from P7 to P14 (Fig. 2). At P7, the periosteal artery invaded into the epiphysis. At this time, PDPN-expressing cells were mainly observed within the periosteum, and some were detected within the vascular tip penetrating the epiphyseal cartilaginous anlage (Fig. 2a). SOC formation and vascularization was initiated at P9. During this process, PDPN-expressing stromal cells began to associate with vascular endothelial cells (Fig. 2b). The SOC became highly vascularized from P10 to P14. PDPN-expressing stromal cells were widely infiltrated in the SOC and were present in the primitive vasculature (Fig. 2c-g). Moreover, during SOC formation, the proliferative expansion of these cells occurred at the penetrating tip of the periosteum (Fig. S1). These findings reveal that epiphyseal PDPN-expressing stromal cells originate from the periosteal cellular component, populate the epiphyseal SOC, and act as pericytes.

**Figure. 2.**
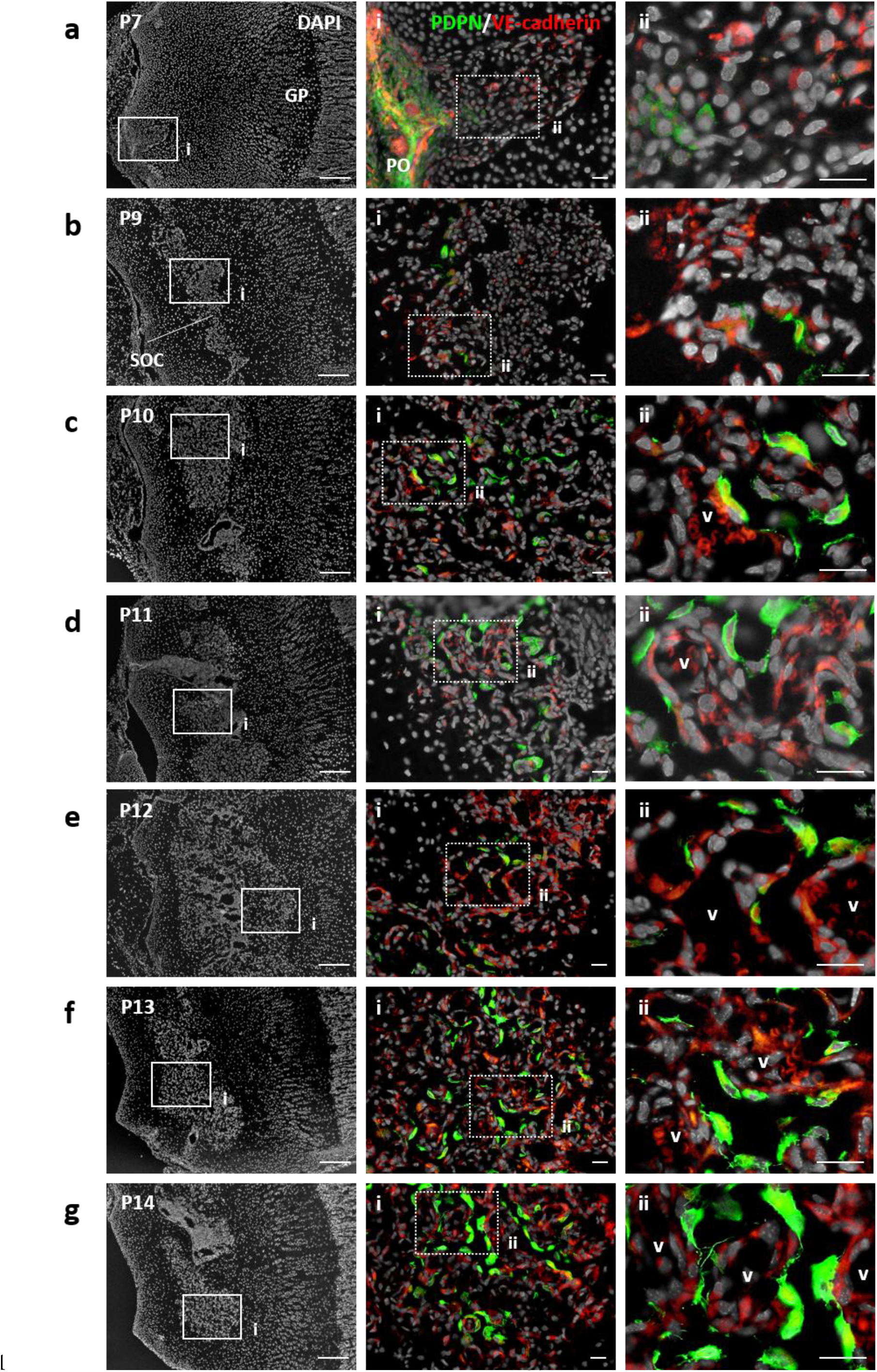
Epiphyseal PDPN-expressing stromal cells originate from the periosteal cellular component and populate the secondary ossification center as pericytes. (A-G) Representative IHC images of postnatal mouse epiphysis at P7 (A), P9 (B), P10 (C), P11 (D), P12 (E), P13(F), and P14 (G). Epiphysis cryo-sections were stained with PDPN, VE-cadherin, and DAPI. Scale bars in the lefts indicate 200 μm. Scale bars in the middle and right panels indicate 50 μm. PDPN: podoplanin, VE-cadherin: vascular endothelial-cadherin, and DAPI: 4′,6-diamidino-2-phenylindole, P7, P9, P10, P11, P12, P13, P14: postnatal day 7, 9, 10, 11, 12, 13, 14

### Epiphyseal PDPN-expressing stromal cells are progeny cells of the SSC lineage

To clarify the cellular characteristics of epiphyseal PDPN-expressing stromal cells, we investigated their potential to act as SSCs. Chan et al.^25^ established a flowcytometric strategy to fractionate SSCs [Lin(-)CD31(-)CD45(-)CD51(+)CD90(-)CD249(-)CD200(+)CD105(-)] and SSC-lineage progenitors [pre-bone cartilage stroma progenitors (preBCSPs), Lin(-)CD31(-)CD45(-)CD51(+)CD90(-)CD249(-)CD200(-)CD105(-)]. We used this strategy to investigate whether epiphyseal PDPN-expressing stromal cells are detected in the SSC or preBCSP fractions. The flowcytometric analysis detected a minor population of epiphyseal PDPN-expressing stromal cells in SSCs and preBCSPs (Fig. 3a and b). The percentages of PDPN-expressing stromal cells in each fraction were 0.202 ± 0.020% and 2.189 ± 0.309% in the SSCs and preBCSPs, respectively (Fig. 3c). The colony formation assay revealed that the clonogenicity of the epiphyseal PDPN-expressing stromal cells was significantly lower than that of primary SSCs and preBCSPs (Fig. 3d). *In vitro* osteo/adipo/chondrogenic differentiation assays showed that epiphyseal PDPN-expressing stromal cells had the potential to differentiate into adipocytes and chondrocytes, but not osteoblasts (Fig. 3e). These findings suggest that epiphyseal PDPN-expressing stromal cells partially exhibit an SSC-lineage phenotype.

**Figure. 3.**
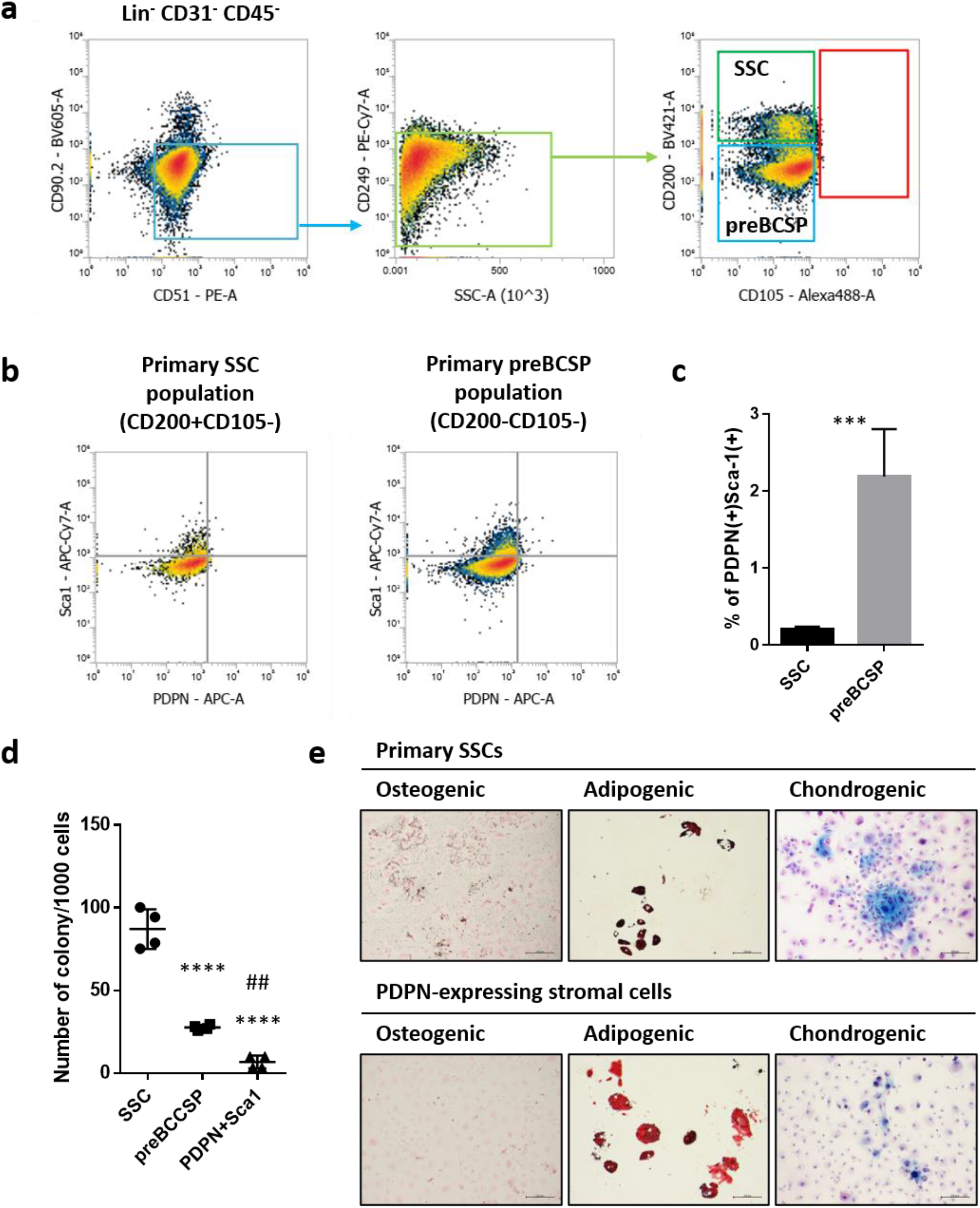
Epiphyseal PDPN-expressing stromal cells partially exhibit the SSC lineage phenotype. (A) Flow cytometric gating strategy for the mouse epiphyseal skeletal stem cell lineage. Cells that were Lin(-)CD31(-)CD45(-)CD51(+)CD90(-)CD249(-)CD200(+)CD105(-) were identified as SSCs. Cells that were Lin(-)CD31(-)CD45(-)CD51(+)CD90(-)CD249(-)CD200(-)CD105(-) were identified as SSC-lineage progenitor and preBCSP populations. (B) Representative flowcytometric scattergrams detecting the PDPN-expressing stromal cells in the epiphyseal primary SSC and preBCSP populations. (C) Quantitative data of the PDPN-expressing stromal cells in the epiphyseal primary SSC and preBCSP populations. ***p < 0.001, as detected by the Student’s *t*-test (n = 4 per group); error bars represent SDMs. (D) Colony formation assay analyzing the clonogenicity of epiphyseal PDPN-expressing stromal cells. Primary SSCs, preBCSPs, and PDPN-expressing stromal cells in the epiphysis were isolated using the cell sorter, and 1000 cells/well were seeded into a 24-well plate. Colonies were counted via Giemsa staining. ****p < 0.0001 v.s. SSCs. ##p < 0.01 v.s. preBCSPs. Statistical analysis was performed using one-way ANOVA with Tukey’s multiple comparison test (n = 4 per group). The error bars represent SDMs. (E) Osteo/adipo/chondrogenic differentiation ability of epiphyseal PDPN-expressing stromal cells. The differentiation of osteogenic, adipogenic, and chondrogenic lineages was evaluated via Von Kossa staining (black), Oil-red staining (red), and Alcian blue staining (sky-blue), respectively. Scale bars indicate 200 μm. SSC: skeletal stem cell, preBCSP: pre-bone cartilage stroma progenitor, PDPN: podoplanin, ANOVA: analysis of variance, SDMs: mean ± standard deviation values of the mean

To confirm their cell lineage, we cultured the primary SSCs with MesenCult and observed the PDPN expression levels. In this experiment, primary SSCs were isolated from the epiphysis at P21, which probably contained P-SSCs as a major fraction and GP-SSCs as a minor fraction. During their culture with MesenCult, epiphyseal primary SSCs differentiated into skeletal lineage-progenitors, including preBCSPs (CD51+CD90-CD249-CD200-CD105-), BCSPs (CD51+CD90-CD249-CD200-CD105+), osteo/chondrogenic progenitors (OC progenitors, CD51+CD90+), and stromal cells (CD51+CD90-CD249+) (Fig. 4a). These *in vitro* differentiated SSC progenies expressed PDPN and Sca-1 at high levels (Fig. 4b). To characterize the PDPN-expressing SSC progenies *in vitro*, we further investigated their surface markers and compared them with those of epiphyseal PDPN-expressing stromal cells. Immunocytochemistry (ICC) showed that PDPN-expressing SSC progenies expressed PDGFRβ and NG2, but not αSMA (Fig. 4c-e), and exhibited a surface marker pattern observed in epiphyseal PDPN-expressing stromal cells (Fig. 1d-f). These observations suggest that epiphyseal PDPN-expressing stromal cells were derived from SSCs.

**Figure. 4.**
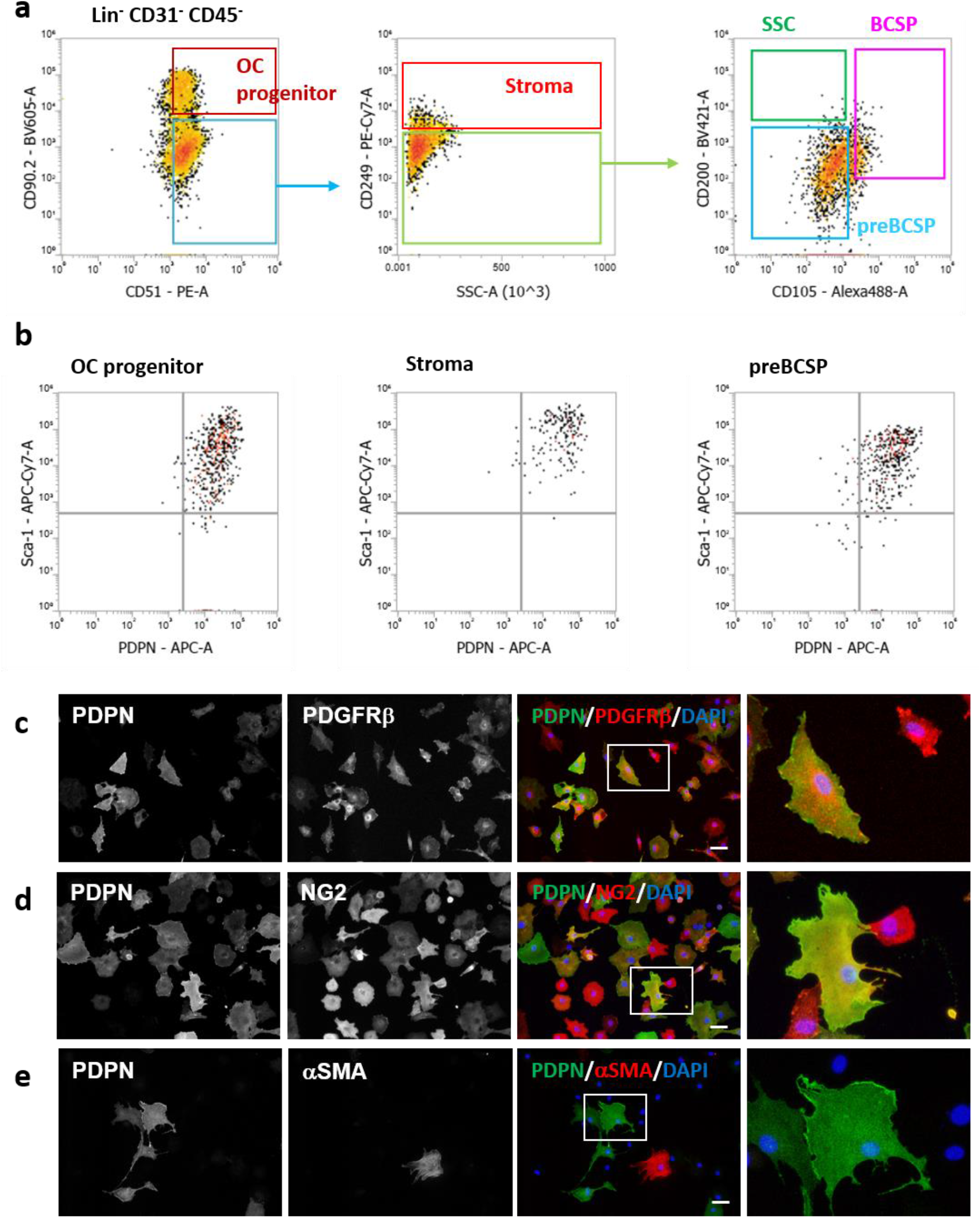
SSCs generate PDPN-expressing progenies with a phenotype identical to that of epiphyseal PDPN-expressing stromal cells. (A) Representative flowcytometric data of *in vitro* SSC progenies generated from primary isolated SSCs during culture with MesenCult. (B) PDPN and Sca-1 expression in *in vitro* generated SSC progenies. (C-E) Representative ICC images of the *in vitro* PDPN-expressing SSC progenies stained with PDPN/PDGFRβ/DAPI (C), PDPN/NG2/DAPI (D), and PDPN/αSMA/DAPI (E). Scale bars in middle panels indicate 50 μm. PDPN: podoplanin, Sca-1: stem cell antigen-1, ICC: immunocytochemistry, DAPI: 4′,6-diamidino-2-phenylindole, PDGFRβ: platelet-derived growth factor receptor-β, NG2: neuron-glial antigen-2, αSMA: α-smooth muscle actin

### PDPN-expressing SSC progenies maintain the HUVEC lumens via the release of angiogenic factors in the xenovascular model

Because epiphyseal PDPN-expressing stromal cells acted as pericytes in the SOC *in vivo*, we hypothesized that these cells could regulate vascular integrity and/or marrow homeostasis. To verify this hypothesis, we used *in vitro* PDPN-expressing SSC progenies as a cellular model of *in vivo* epiphyseal PDPN-expressing stromal cells. In addition, we established a xenovascular model by co-culturing Human umbilical vein endothelial cells (HUVECs) with *in vitro* PDPN-expressing SSC progenies. In the xenovascular model, non-endothelial cells with fibroblastic morphologies were attached onto HUVEC vascular-like cords (Fig. 5a). ICC revealed that these pericyte-like cells expressed PDPN, PDGFRβ, and NG2, but not VE-cadherin (Fig. 5b-d). These observations indicate that *in vitro* PDPN-expressing SSC progenies acted as pericytes surrounding the HUVEC cords, and mimicked epiphyseal PDPN-expressing stromal cells present on the primitive vascular beds in the SOC. Next, we investigated whether SSC progenies expressing PDPN *in vitro* maintained HUVEC lumens. Compared to the conditions used for the monoculture of HUVECs, the parameters used to evaluate HUVEC lumen integrity (the number of junctions, the number of segments, the number of meshes, and the total mesh area) were significantly maintained in the xenovascular-model. (Fig. 5e-i). Moreover, the luminal regulation of PDPN-expressing SSC progenies was sustained for at least 6 days (Fig. S2). These observations indicate that PDPN-expressing SSC progenies consolidate the HUVEC lumens *in vitro*.

**Figure. 5.**
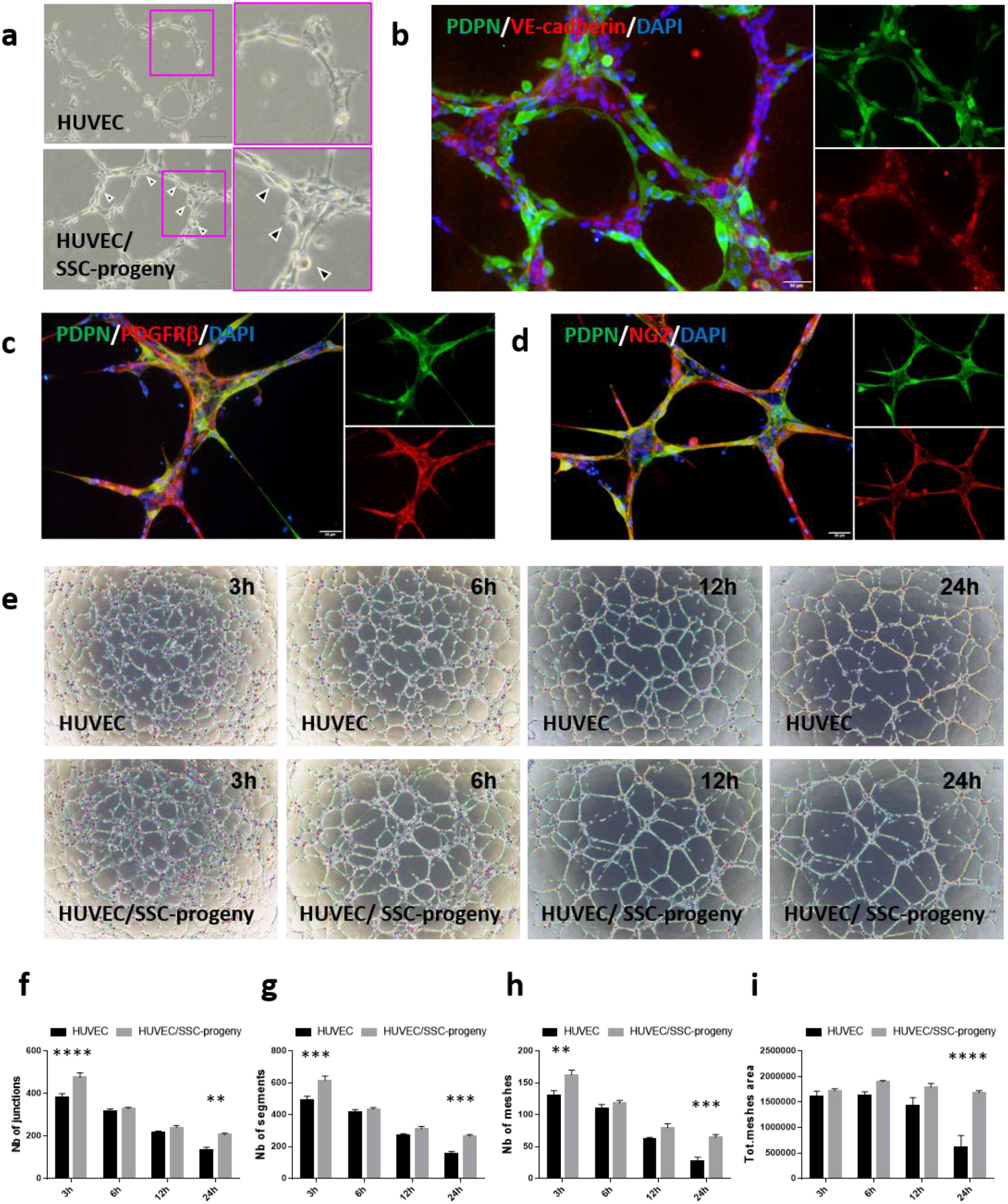
*In vitro* PDPN-expressing SSC progenies consolidate HUVEC capillary-like lumens in a xenovascular model. (A) Representative optical microscope images of the vascular-like lumen of HUVECs (upper panel) and the xenovascular model co-cultured with HUVECs and PDPN-expressing SSC progenies *in vitro* (lower panel). Arrow heads indicate non-endothelial cells with fibroblastic morphologies attached onto HUVEC cords. (B) Representative ICC images of the xenovascular model stained with PDPN, VE-cadherin, and DAPI. Scale bar indicates 50 μm. (C and D) Representative ICC images of the xenovascular images stained with PDPN/PDGFRβ/DAPI (C) and PDPN/NG2/DAPI (D). Scale bars indicate 50 μm. (E) Time series images of HUVEC vascular-like lumens (upper panels) and the xenovascular models co-cultured with HUVECs and PDPN-expressing SSC progenies *in vitro* (lower panels). The time displayed on each image indicates the time point at the start of the culture process. (F-I) Quantitative analysis of vascular lumen integrity in the xenovascular model. The parameters used to evaluate lumen vascularization, including the number of junctions (F), the number of segments (G), the number of meshes (H), and the total mesh area (I), were measured using the angiogenesis analyzer tool. **p < 0.01. ***p < 0.001. ****p < 0.0001. Statistical analysis was performed via two-way ANOVA and Sidak’s multiple comparison test (n = 5 per group). The error bars represent SEMs. HUVEC: human umbilical vein endothelial cell, PDPN: podoplanin, SSC: skeletal stem cell, ICC: immunocytochemistry, VE-cadherin: vascular endothelial-cadherin, DAPI: 4′,6-diamidino-2-phenylindole, PDGFRβ: platelet-derived growth factor receptor-β, NG2: neuron-glial antigen-2, ANOVA: analysis of variance, SEMs: mean ± standard error values of the mean

To investigate the mechanism of vascular regulation by *in vitro* PDPN-expressing SSC progenies, we analyzed soluble factors in conditioned medium derived from PDPN-expressing SSC progenies (SSC-progeny CM). The HUVEC proliferation assay showed that SSC-progeny CM significantly accelerated cell proliferation (Fig. 6a). The proliferative activity was much higher than that of the commercially available endothelial cell growing medium (EGM2). Scratch assays revealed that SSC-progeny CM also facilitated HUVEC migration (Fig. 6b). Therefore, we investigated whether SSC-progeny CM consolidated the HUVEC lumens. When compared to the non-conditioned medium (Non-CM), the SSC-progeny CM significantly enabled the parameters required for HUVEC lumen integrity to be maintained (Fig. 6c-g); this mimicked the behavior in the xenovascular model containing HUVECs and PDPN-expressing SSC progenies (Fig. 5e-i). To profile the soluble factors regulating HUVEC lumen integrity, we performed a protein array analysis of 53 angiogenesis-related factors. The results revealed that SSC-progeny CM contained various angiogenic factors, and the spot intensities of 7 angiogenic factors, i.e., IGFBP-2^33^, osteopontin^34^, CCL2^35^, MMP-3^36, 37^, CXCL12^38^, PAI-1^39–41^, and TSP-2^42, 43^ were particularly increased (Fig. 6h and Fig. S3). These data indicate that PDPN-expressing SSC progenies autonomously secrete various angiogenic factors that maintain HUVEC lumens in concert. Thus, we suggest that epiphyseal PDPN-expressing stromal cells positively regulate the integrity of the primitive vasculature, via the release of angiogenic factors.

**Figure. 6.**
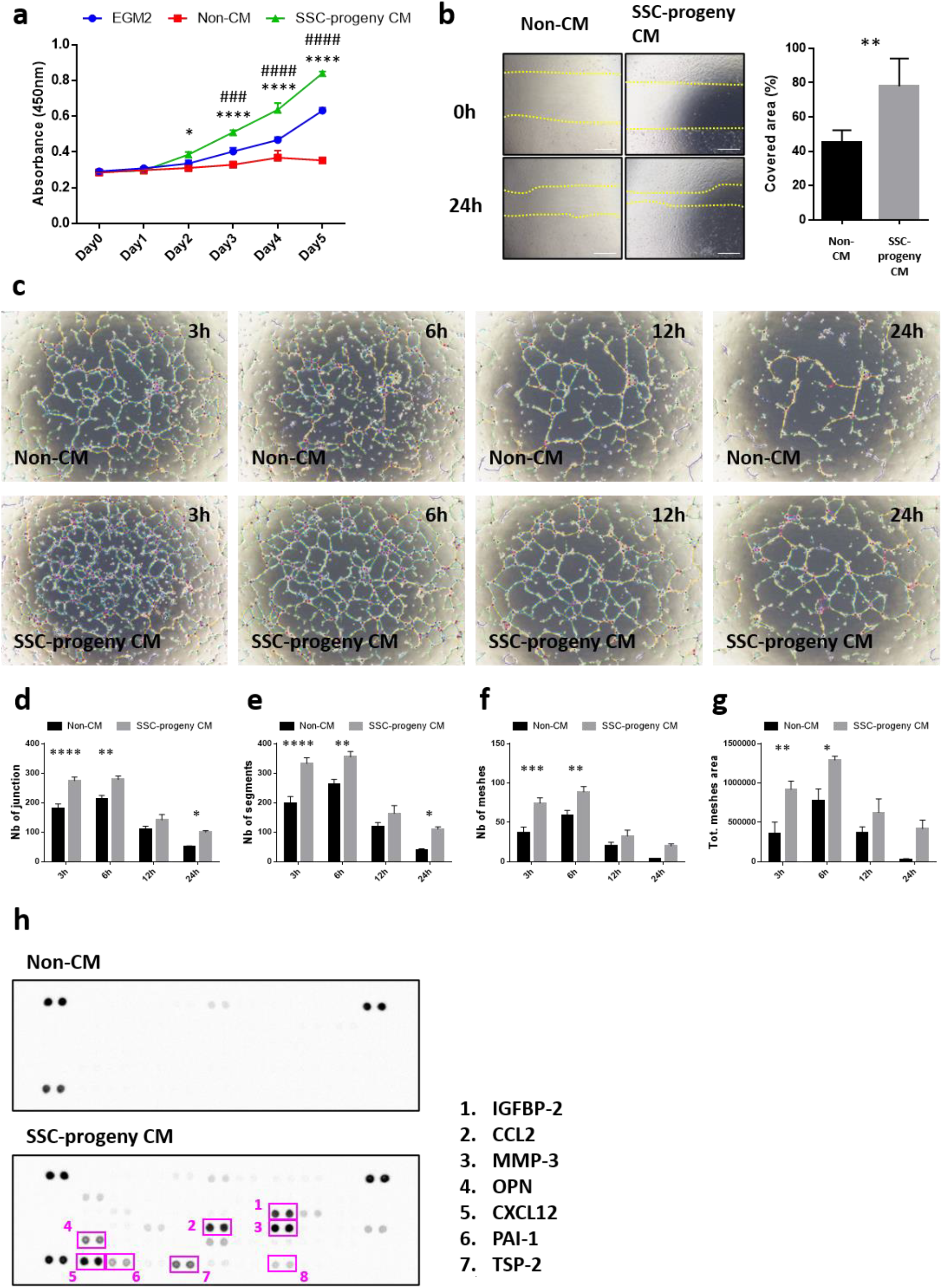
PDPN-expressing SSC progenies autonomously release various angiogenic factors that coordinate with each other to maintain HUVEC lumens *in vitro*. (A) HUVEC proliferation assay. HUVECs were cultured with the EGM2, non-CM, and SSC-progeny CM media. HUVEC proliferation was assessed via the WST-8 assay. *p < 0.05 v.s. Non-CM. ****p < 0.0001 v.s. Non-CM. ###p < 0.001 v.s. EGM2. ####p < 0.0001 v.s. EGM2. Statistical analysis was performed using two-way ANOVA and Sidak’s multiple comparison test (n = 3 per group). The error bars represent SEMs. (B) HUVEC scratch assay. The panel on the left indicates the representative optical microscopic images of scratched HUVEC monolayers cultured with non-CM and SSC-progeny CM media at 0 h or 24 h. The panel on the right indicates the quantitative data of the covered area, 24 h after culturing cells with non-CM and SSC-progeny CM media. **p < 0.01, as detected by the Student’s t-test (n = 5 per group). Scale bars indicate 200 μm. (C) Time series images of HUVEC vascular-like lumens in non-CM (upper panels) and SSC-progeny CM (lower panels) media. (D-G) Quantitative analysis of HUVEC vascular lumen integrity using non-CM or SSC-progeny CM media. The parameters used for evaluating lumen vascularization, including the number of junctions (D), the number of segments (E), the number of meshes (F), and the total mesh area (G), were measured using the angiogenesis analyzer tool. *p < 0.05. **p < 0.01. ***p < 0.001. ****p < 0.0001. Statistical analysis was performed by two-way ANOVA and Sidak’s multiple comparison test (n = 5 per group). The error bars represent SEMs. (H) Profiling the soluble factors to regulate HUVEC lumen integrity in the SSC-progeny CM medium. The soluble factors were profiled by using the Proteome Profiler Mouse Angiogenesis Array Kit. The top and bottom panels indicate the array images generated with non-CM and SSC-progeny CM media, respectively. PDPN: podoplanin, SSC: skeletal stem cell, HUVEC: human umbilical vein endothelial cell, EGM2: endothelial growth medium-2, non-CM: EGM2-basal medium supplemented with non-conditioned medium, SSC-progeny CM: EGM2-basal medium supplemented with *in vitro* PDPN-expressing SSC-progeny conditioned medium

### PDPN-expressing SSC progenies secrete components of the basement membrane matrices in response to Notch-mediated interactions with HUVECs

Vascular integrity is not only maintained by endothelial cell survival, but also by the microenvironments in the perivascular space. ECM is the one of major environmental factors secreted by endothelial cells and their pericytes that maintains vascular homeostasis^44, 45^. We therefore investigated whether *in vitro* PDPN-expressing SSC progenies secrete collagenous and non-collagenous ECM components. First, we determined the expression of type IV collagen and laminin isoforms in *in vitro* PDPN-expressing SSC progenies under monoculture conditions (Fig. 7a and b). The *in vitro* PDPN-expressing SSC progenies endogenously produce type IV collagen and laminin; however, the extracellular deposition of these ECMs was not observed. Second, we analyzed the extracellular deposition of basement membrane ECMs in the xenovascular model (Fig. 7c-f). The extensive deposition of type IV collagen and laminin isoforms was observed in close proximity to the luminal cords (Fig. 7c and e). Quantitative image analysis showed that the levels of the deposited type IV collagen and laminin isoforms were significantly increased during the co-culture of HUVECs with *in vitro* PDPN-expressing SSC progenies, compared to the levels observed in HUVEC monoculture (Fig. 7d and f). These findings indicate that interactions with HUVECs induce *in vitro* PDPN-expressing SSC progenies to secrete ECMs in the periluminal space.

**Figure. 7.**
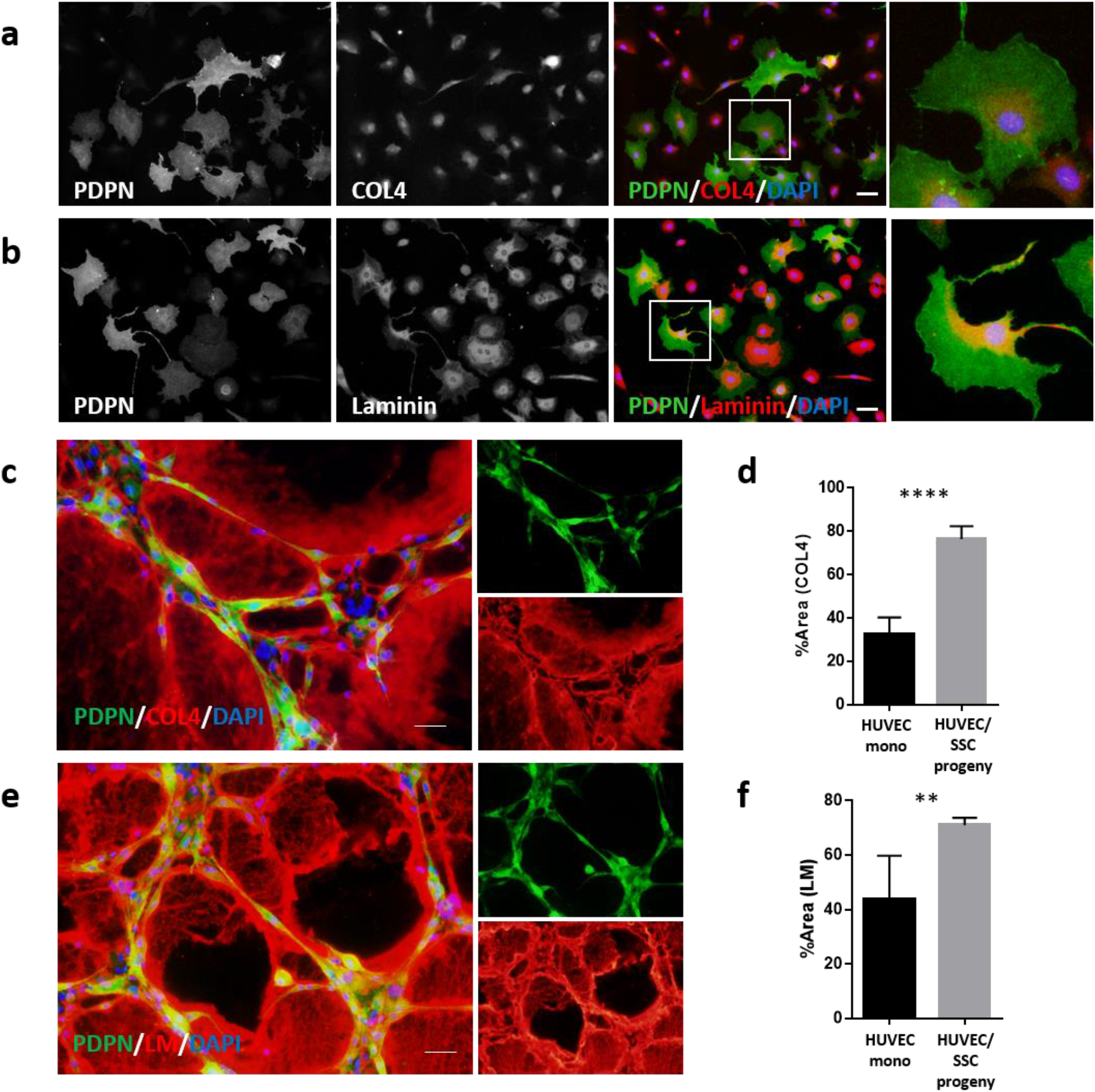
Cell-cell interactions with HUVECs induce PDPN-expressing SSC progenies to release ECMs in the periluminal space *in vitro*. (A-B) Representative ICC images of monocultured PDPN-expressing SSC progenies stained *in vitro* with type IV collagen (A) and laminin isoforms (B). (C-F) Basement membrane ECM deposition at the periluminal space in the xenovascular model. (C and E) Representative ICC images of the xenovascular model co-cultured with HUVECs and PDPN-expressing SSC progenies stained *in vitro* with type IV collagen (C) and laminin isoforms (E). (D and F) Quantitative area analysis of deposited type IV collagen (D) and laminin isoforms (F) in the xenovascular model. **p < 0.01. ****p < 0.0001. Statistical analysis was performed using the Student’s t-test (n = 5 per group). The error bars represent SEMs. Scale bars indicate 50 μm. ICC: immunocytochemistry, PDPN: podoplanin, SSC: skeletal stem cell, HUVECs: human umbilical vein endothelial cells

Next, we examined whether the cell-cell interactions with HUVECs altered the ECM transcript pattern of PDPN-expressing SSC progenies. Xenovascular capillaries were enzymatically dissociated, and co-cultured PDPN-expressing SSC progenies were isolated *in vitro* via cell sorting (Fig. 8a). After immunostaining with anti-mouse PDPN-APC and anti-human CD31-FITC antibodies, PDPN-expressing SSC progenies were identified to be mouse PDPN-positive/human CD31-negative cells (Fig. 8b). RT-qPCR revealed that the transcript expression level of mouse vascular basement membrane ECM-related genes, i.e., type IV collagen alpha-chains (*Col4a1* and *Col4a2*) in PDPN-expressing SSC progenies was significantly upregulated upon co-culture with HUVECs (Fig. 8c). We evaluated the expression of genes related to non-collagenous basement membrane ECMs, such as laminin alpha-chains (*lama4* and *lama5*) and nidogen isoforms (*Nid1* and *Nid2*) (Fig. 8d). Among these non-collagenous ECM-related genes, the expression of *lama5* and *Nid1* was significantly upregulated in PDPN-expressing SSC progenies upon co-culture with HUVECs, whereas the expression of *Lama4* was not altered and that of *Nid2* was significantly decreased. However, the expression of genes encoding non-basement membrane fibrillar collagen, i.e., *Col1a1* and *Col3a1*, was downregulated in PDPN-expressing SSC progenies and remained unaltered after co-culture with HUVECs (Fig. 8e). These findings show that the interaction of PDPN-expressing SSC progenies with HUVECs results in the switching of the ECM-phenotype of PDPN-expressing SSC progenies to the basement-membrane‒dominant state.

**Figure. 8.**
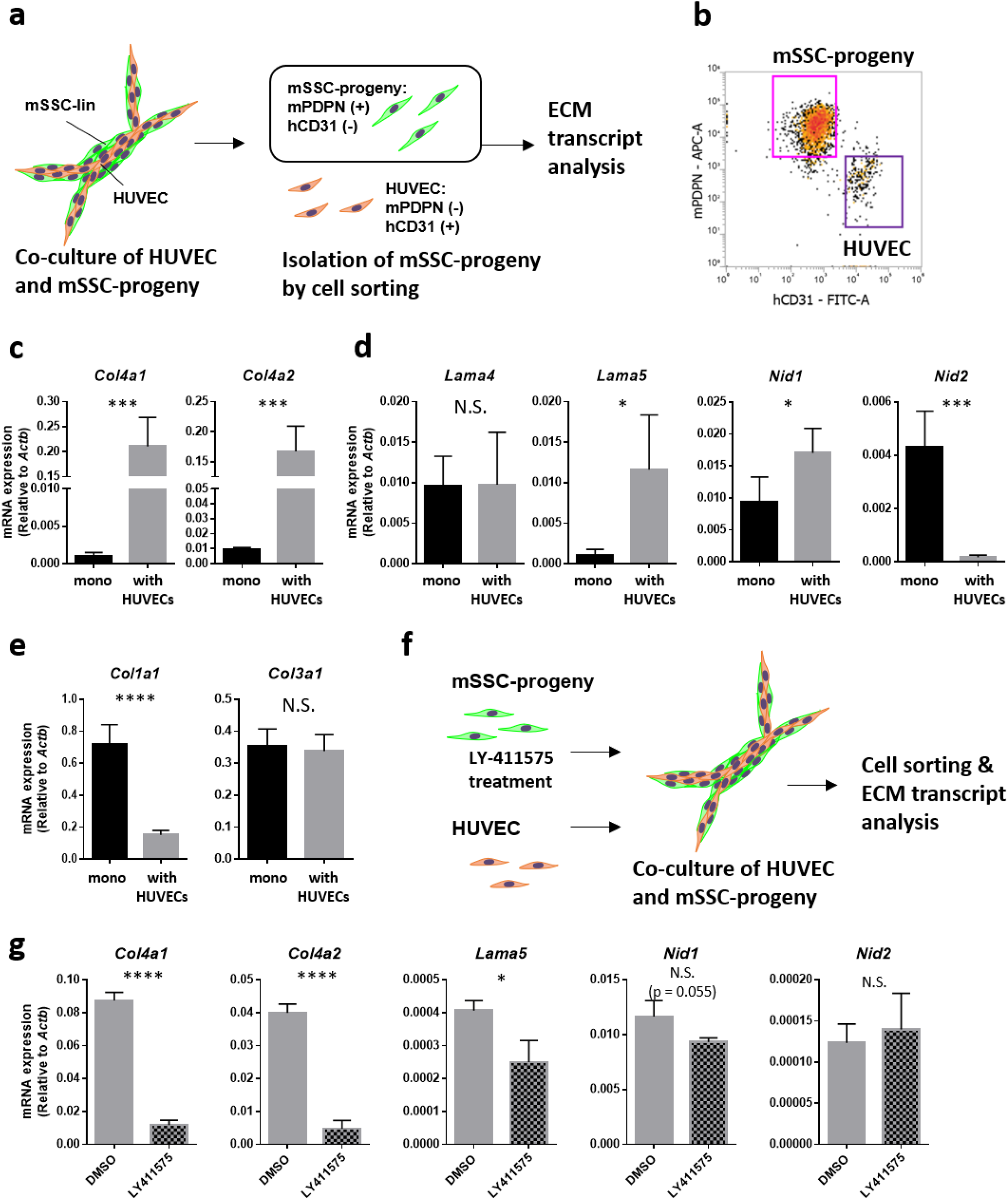
Notch-associated cell-cell interactions with HUVECs upregulate basement membrane ECM expression of PDPN-expressing SSC progenies *in vitro*. (A) Strategy for isolating co-cultured PDPN-expressing SSC progenies in the xenovascular model. (B) Representative flow cytometric scattergram detecting HUVECs and PDPN-expressing SSC progenies via the staining of human CD31 and mouse PDPN. Distinctively separated mouse PDPN-expressing SSC progenies were isolated using the cell sorter. (C) RT-qPCR evaluating the expression of the genes encoding proteins present in collagenous vascular basement membrane ECMs, such as type IV collagen alpha-chains (*Col4a1* and *Col4a2*). (D) RT-qPCR evaluating the expression of genes encoding the proteins present in the non-collagenous vascular basement membrane ECMs, such as laminin alpha-chains (*lama4* and *lama5*) and nidogen isoforms (*Nid1* and *Nid2*). (E) RT-qPCR evaluating the expression of genes encoding the proteins present in non-vascular basement membrane ECMs, such as fibrillar type I and type III collagen (*Col1a1* and*Col3a1*). (F) Strategy for experiments involving the xenovascular model and LY-411575, a Notch pathway inhibitor. Prior to their co-culture with HUVECs, PDPN-expressing SSC progenies were pre-treated with LY-411575 *in vitro*. Isolated PDPN-expressing SSC progenies were subjected to ECM transcript analysis. (G) LY-411575 suppresses the basement membrane ECM up-regulation of PDPN-expressing SSC progenies during cell-cell interactions with HUVECs. In this experiment, we targeted the ECM genes *Col4a1*, *Col4a2*, *Lama5*, *Nid1*, and *Nid2*, which were altered during co-culture with HUVECs. *p < 0.01. **p < 0.01. ***p < 0.01. ****p < 0.0001. N.S. indicates non-significance differences. Statistical analysis was performed by the Student’s t-test (n = 5 per group). The error bars represent SEMs. PDPN: podoplanin, SSC: skeletal stem cell, HUVECs: human umbilical vein endothelial cells, ECM: extracellular matrix

Notch activation reportedly upregulated the expression of basement membrane ECM-related genes in BM-SSCs *in vivo*^46^. Therefore, we investigated whether Notch signals were responsible for alterations in the ECM phenotype of PDPN-expressing SSC progenies *in vitro*. After treating SSC progenies with LY-411575, a Notch pathway inhibitor in the background of co-culture with HUVECs, ECM expression levels were evaluated (Fig. 8f). We first determined the effects of LY-411575 on the formation of vascular-like lumens in the xenovascular model (Fig. S4). LY-411575 did not affect the morphology and vascular integrity of the xenovascular model. Next, we investigated whether LY-411575 altered the expression of ECM-related genes in SSC progenies co-cultured with HUVECs (Fig. 8c and d). Notch pathway inhibition significantly suppressed the HUVEC-induced upregulation of *Col4a1*, *Col4a2*, *Lama5*, and *Nid1* (Fig. 8g). LY-411575 did not affect *Nid2* expression in the background of HUVEC co-culture. These data reveal that Notch-mediated interactions of PDPN-expressing SSC progenies with HUVECs result in the switching of the ECM phenotype of SSC progenies to the basement membrane-dominant state.

## Discussion

Bone marrow PDPN-expressing stromal cells generate megakaryopoietic and erythropoietic microenvironments in the perivascular space of the bone marrow in adult mice^28, 29^. However, their contribution to marrow development and homeostasis has been unclear. In this study, we found that PDPN-positive periosteal cells infiltrated the cartilaginous anlage of the postnatal epiphysis and acted as pericytes in the primitive SOC vasculature (Fig. S5a). In addition, we revealed that PDPN-expressing stromal cells originate from SSCs. Based on the findings obtained using the *in vitro* xenovascular model, we propose that PDPN-expressing stromal cells maintain vascular integrity via the release of angiogenic factors and vascular basement membrane ECM related molecules (Fig. S5b). In addition, Notch-mediated interactions of PDPN-expressing stromal cells with endothelial cells induce the switching of the phenotype associated with the ECM expression pattern to the pericyte phenotype.

PDPN is a mucin-type transmembrane protein that binds to C-type lectin-like receptor-2 (CLEC-2, also known as CLEC1B) expressed on platelets and megakaryocytes^47–49^. In non-bone marrow tissues, PDPN is expressed by multiple cell types^50^, such as type I alveolar epithelial cells (ACE1)^51^ and lymphatic endothelial cells (LECs)^52^. The interaction between PDPN on ACE1 and CLEC-2 on platelets regulates neonatal lung development^53^. The interaction of PDPN on LECs and CLEC-2 on platelets promotes lymphatic vessel development in the embryos^54, 55^. In the lymph node, fibroblastic reticular cells (FRCs) express PDPN and maintain lymph node homeostasis by regulating the integrity of high endothelial venules (HEVs)^56, 57^. During lymph node hemorrhage (e.g., increased lymphocyte trafficking such as chronic inflammation), FRC PDPN interacts with CLEC-2 on extravasated platelets. The platelets activated via the PDPN/CLEC-2 axis locally release sphingosine-1-phosphate in the perivenular space, resulting in increased HEV integrity^56^. The PDPN/CLEC-2 axis is a key determinant of vascular integrity that maintains vascularized tissue homeostasis and development. Bone marrow PDPN-expressing stromal cells are considered to contribute to nascent marrow homeostasis by consolidating vascular integrity in the same manner as the PDPN/CLEC-2 axis in the lymph node.

ECM molecules play an important role in the formation of the vasculature and maintenance of its integrity. In the vascular system, the ECM forms two types of structures, i.e., the interstitial matrix and the basement membrane^44^. The basement membrane is a sheet-like structure composed of type IV collagen, laminins (laminin-411 and laminin-511), nidogens, and perlecan, which represents a physiological barrier to the movement of intra/extravascular soluble molecules and migrating cells^45, 58^. In addition, the basement membrane provides a scaffold that supports vascular lumen formation and the interaction between the endothelium and pericytes^59^. The use of *in vitro* PDPN-expressing SSC progenies in our xenovascular model suggests that PDPN-expressing stromal cells prime the perivascular environment via the secretion of basement membrane ECM molecules (Fig. 7 c-f). Further, we observed that PDPN-expressing SSC progenies switched their ECM expression pattern, which resulted in the dominance of basement membrane components, via Notch-mediated interactions with endothelial cells (Fig. 8c-g). Endothelial cells express Notch ligands, such as delta-like 4 and jagged-1, which activate the Notch pathway in pericytes or perivascular cells via cell-cell interactions^60–64^. Knockout of Notch pathway intermediaries results in vascular defects or pericyte dysfunction^65–69^. This evidence indicates the important role played by Notch signaling in vascular development. We hypothesize that bone marrow PDPN-expressing stromal cells switch their cellular phenotype to the pericyte phenotype via Notch-mediated interactions with the endothelium, and this process promotes marrow vascularization.

In this study, we have shown the vascular regulatory functions of epiphyseal marrow PDPN-expressing stromal cells using the *in vitro* xenovascular model. However, this study has a few limitations. To further demonstrate their role in bone marrow physiology including disease pathophysiology, an *in vivo* cell-fate reporter or depletion model that specifically targets the marrow PDPN-expressing stromal cells must be established. These *in vivo* models would demonstrate the detailed mechanism by which PDPN-expressing stromal cells regulate bone marrow development and homeostasis. Furthermore, it must be examined whether the marrow PDPN-expressing stromal cells are involved in bone marrow development during embryogenesis (especially POC-associated marrow development). These questions need to be addressed in future studies.

Our study offers a new perspective in understating how stromal cells regulate nascent bone marrow development and homeostasis. This study can be used as a basis for further studies to comprehensively examine the contribution of stromal cells to bone marrow developmental physiology. This would provide insights into the mechanism of the bone and bone marrow development

## Online Methods

### Mice

C57BL/6NcrSlc mice were purchased from CLEA Japan, Inc (Tokyo, Japan). They were bred and maintained under standard conditions [a 12 hour light/dark cycle with stable temperature (25 °C) and humidity (60%)]; the mice that were 7–21 postnatal days old were selected for the experiments. This study was approved by the animal care and use committee at the Nagoya University Faculty of Medicine, Nagoya, Japan (D210596-003).

### Flow cytometry

Femurs were harvested from mice at P21. To obtain epiphyseal marrow stromal cells, dissected epiphyses were gently smashed using a mortar and further cut to small pieces. After a few washes with ice-cold PBS containing 10% FBS (Sigma-Aldrich, Tokyo, Japan), the epiphyseal pieces were digested with 0.2% (w/v) collagenase (Wako, Tokyo, Japan) for 2 h at 37 °C and agitated at 100 rpm. After collagenase digestion, the epiphyseal pieces were further crushed in a mortar with ice-cold PBS containing 10% FBS. Harvested cells were passed through a 40-micrometer cell strainer (Corning, Bedford, MA, USA). The cell suspension was centrifuged at 280 × *g* for 5 min, at 4 °C. The cell pellet was hemolyzed with sterilized ultrapure water for 6 s, and washed with ice-cold PBS containing 10% FBS.

To obtain diaphyseal marrow stromal cells, the marrow was flushed from the dissected diaphysis using a 21-gauge needle (Terumo, Tokyo, Japan). The flushed diaphyseal marrow was suspended in ice-cold PBS containing 10% FBS and passed through a 40-micrometer cell strainer. After centrifugation, the cell pellet was resuspended and hemolyzed using the ACK buffer (155 mM NH_4_Cl, 10mM KHCO_3_, 0.1mM EDTA). The hemolyzed cells were probed with antibodies against hematopoietic lineage markers (CD4, CD8, B220, TER-119, Ly-6G, CD11b, F4/80, and CD71). Cells of the hematopoietic lineage were depleted using sheep anti-rat immunoglobulin G (IgG) polyclonal antibody (pAb)-conjugated magnetic beads (Dynabeads, Thermo Fisher Scientific, Waltham, MA, USA). Cells that were not of hematopoietic lineage were harvested and washed with ice-cold PBS containing 10% FBS. For enrichment of PDPN-positive cells, epiphyseal or diaphyseal stromal cells were probed using the anti-PDPN APC conjugate (Clone: 8.1.1, Biolegend, San Diego, CA, USA) and isolated as APC-positive fraction using anti-APC Microbeads (Miltenyi Biotec, San Jose, CA, USA) on an LS column (Miltenyi Biotec). The depletion antibodies used in the study are listed in Table S1.

Harvested cells were probed with fluorescence-conjugated antibodies or their isotype controls. Flow cytometry was performed using a three-laser Attune NxT (Ex.405/488/637 nm, Thermo Fisher Scientific). Cell sorting was performed using a four-laser FACS Aria II (Ex.355/407/488/633 nm, BD Bioscience, San Jose, CA, USA). The fluorescence-conjugated antibodies used in this study are listed in Table S1.

### Immunohistochemistry

Femurs were fixed with 4% paraformaldehyde (PFA, Wako, Tokyo, Japan) for 24 h and subsequently decalcified for 16 h using K-CX (FALMA, Tokyo, Japan). After washing with diluted water, bone pieces were incubated in 30% sucrose (Wako, Tokyo, Japan) for cryoprotection. Treated tissues were embedded in Tissue-Tek O.C.T. Compound (Tissue-Tek, Sakura, Japan) at -80°C. Frozen sections were sectioned to generate 10-micrometer-thick sections and blocked with phosphate-buffered saline (PBS) containing 3% bovine serum albumin (Sigma-Aldrich, Tokyo, Japan) and 2% goat serum (Sigma-Aldrich). Sections were probed overnight with primary antibodies against PDPN, VE-cadherin, NG2, and αSMA—diluted with the blocking reagent—at 4 °C. Then, the sections were probed with secondary antibody conjugates, including anti-Syrian hamster IgG Alexa 488 conjugate (for PDPN, A21110, Thermo Fisher Scientific, 1:1000 diluted) and anti-rabbit IgG Alexa 568 conjugate (for VE-cadherin, PDGFRβ, and NG2, A11034, Thermo Fisher Scientific, 1:2000 diluted), and anti-mouse IgG Alexa 568 conjugate (for αSMA, A11004, Thermo Fisher Scientific, 1:2000 diluted) for 1.5 h at 25°C. The sections were mounted with VECTASHIELD Antifade Mounting Medium containing DAPI (Vector Laboratories, Burlingame, CA, USA), and observed under an upright fluorescence microscope (AX80, Olympus, Tokyo, Japan). The primary antibodies used in the study are listed in Table S1.

### Immunocytochemistry

Cells were seeded onto 15-millimeter Fisherbrand™ Coverglass for Growth™ Cover Glasses (Thermo Fischer Scientific) and cultured in 12-well culture plates at 37°C, 5% CO_2_. HUVECs were passaged 4 to 8 times, whereas primary SSCs were passaged <3 times. The cells were fixed with 4% PFA. To achieve permeabilization, cells were incubated with 0.1% Triton X-100 (Wako, Tokyo, Japan) prepared in PBS for 10 min. After washing with PBS, the cells were blocked with PBS containing 3% bovine serum albumin and 2% goat serum for 1 h at 25°C. Cells were probed overnight with primary antibodies against PDPN, VE-cadherin, NG2, αSMA, COL4, and laminins at 4 °C. Then cells were probed with secondary antibody conjugates, including anti-Syrian hamster IgG Alexa 488 conjugate (for PDPN, A21110, Thermo Fisher Scientific, 1:1000 diluted) and anti-rabbit IgG Alexa 568 conjugate (for VE-cadherin, PDGFRβ, NG2, COL4, and laminins, A11034, Thermo Fisher Scientific, 1:2000 diluted), and anti-mouse IgG Alexa 568 conjugate (for αSMA, A11004, Thermo Fisher Scientific, 1:2000 diluted) for 1.5 h at 25°C. Cells were mounted using VECTASHIELD Antifade Mounting Medium containing DAPI (Vector Laboratories, Burlingame, CA, USA), and observed under an inverted fluorescence microscope (IX73, Olympus, Tokyo, Japan). Acquired images were quantitatively analyzed using Image J 1.46r (http://rsb.info.nih.gov/ij/).

### Cell culture

HUVECs (TaKaRa Bio, Shiga, Japan) were cultured using Endothelial Cell media 2 (EGM2, TaKaRa Bio) and penicillin/streptomycin/amphotericin B (Wako, Osaka, Japan). Isolated mouse primary SSCs were cultured using the mouse MesenCult Expansion Kit with L-glutamine (Wako) and penicillin/streptomycin/amphotericin B (Wako).

### Colony formation assay

Isolated cells were seeded onto a 12-well culture plate (1000 cells/well, Thermo Fisher Scientific) and cultured using the Complete MesenCult expansion medium (Stem Cell Technologies, Vancouver, Canada) for 7 days. Cells were stained with Giemsa staining solution (Muto pure chemicals, Tokyo, Japan).

### *In vitro* mesenchymal tri-lineage differentiation

For osteogenic differentiation, cells were seeded into a 24-well plate (4 × 10^5^ cells/cm^2^) and cultured using the Complete MesenCult expansion medium. After 24 h, the culture medium was replaced with the Complete mouse MesenCult Osteogenic Medium (Stem Cell Technologies, Vancouver, Canada) containing L-glutamine (Wako) and penicillin/streptomycin/amphotericin B (Wako), and cells were cultured for 12 days. Osteoblastic differentiation was investigated by evaluating calcium deposition via Von Kossa staining. Briefly, cells were fixed with 4% PFA and washed with PBS. After rinsing cells with distilled water, the deposited calcium was stained using a Calcium Stain Kit (ScyTek laboratories, Utah, USA); cells were subsequently counterstained with the Fast Red solution.

To achieve adipogenic differentiation, cells were seeded in a 24-well plate (1 × 10^5^ cells/cm^2^) and maintained using the Complete MesenCult expansion medium. After 24 h, the culture medium was replaced with the Complete mouse MesenCult Adipogenic Differentiation Medium (Stem Cell Technologies) containing L-glutamine (Wako) and penicillin/streptomycin/amphotericin B (Wako), and cells were cultured for 12 days. Adipogenic differentiation was evaluated by staining adipocytes with Oil Red O (Wako). A working solution of Oil Red O was prepared by mixing Oil Red O stock solution [0.15 g Oil Red O (Wako) in 100% isopropanol (Wako)] and distilled water at a dilution of 6:4. It was filtered after 20 min. Cells were fixed with 4% PFA, washed with PBS, and incubated with 60% isopropanol for 1 min. Then, cells were incubated for 20 min with a working solution of Oil red O at room temperature, rinsed with 60% isopropanol, and washed twice with PBS.

To achieve chondrogenic differentiation, cells were seeded in a 24-well plate (4 × 10^5^ cells/cm^2^) and cultured using the Complete MesenCult expansion medium. After 24 h, the culture medium was replaced with the Complete mouse MesenCult-ACF Chondrogenic Differentiation Medium (Stem Cell Technologies) and penicillin/streptomycin/amphotericin B (Wako), and cells were cultured for 12 days. Chondrogenic differentiation was evaluated by staining chondrocyte-associated mucopolysaccharides with Alcian Blue. Cells were fixed with 4% PFA, washed with PBS, and treated with 3% acetic acid. Then, cells were incubated with an Alcian Blue (pH of 2.5; Muto Pure Chemicals, Tokyo, Japan) for 30 min at room temperature and washed with 3% acetic acid. After rinsing with distilled water, the cells were counterstained with the Fast Red solution for 5 min, and washed twice with distilled water.

### RNA extraction and reverse transcribed-quantitative polymerase chain reaction (RT-qPCR)

Total RNA was extracted by using the ReliaPrep RNA Cell Miniprep System (Promega, Fitchburg, WI, USA). First strand cDNA was synthesized using the PrimeScript II 1st strand cDNA Synthesis Kit (TaKaRa Bio, Shiga, Japan), according to the manufacturer’s instructions. Multiplex qPCR was performed using the TaqMan Gene expression Master Mix (Thermo Fisher Scientific), PrimeTime qPCR Assay (Integrated DNA Technology, Singapore, Republic of Singapore), and Thermal Cycler Dice Real Time System (TaKaRa Bio). The cycling conditions were: 95 °C for 10 min and 40 cycles of denaturation at 95 °C for 15 s and annealing/extension at 60 °C for 1 min. Fluorescence intensity was measured at every annealing/extension step. The qPCR probes used in this study are listed in Table S2.

### HUVEC capillary formation and xenovascular model

HUVECs (0.53 × 10^5^/cm^2^) and/or *in vitro* SSC progenies (0.39 × 10^5^/cm^2^) were seeded onto a cell culture plate or coverglass 8-well chamber (Iwaki, Tokyo, Japan) coated with the Corning® Matrigel® Growth Factor Reduced (GFR) Basement Membrane Matrix (Corning), and cultured with Complete EGM2 supplemented with 2% Matrigel and 10 ng/mL VEGF-A (Miltenyi Biotech). HUVEC lumen integrity was analyzed using the Angiogenesis Analyzer for ImageJ tool (http://image.bio.methods.free.fr/ImageJ/?Angiogenesis-Analyzer-for-ImageJ&lang=en). To evaluate the vascular lumen integrity, we determined several parameters, i.e., the number of junctions, the number of segments (segments are elements delimited by two junctions), the number of meshes (meshes are areas enclosed by segments), and the total mesh area.

For Notch pathway inhibition, *in vitro* SSC progenies were treated with 1 μM LY-411575 (Sigma-Aldrich) for 24 h, and re-seeded into the xenovascular model with HUVECs. To achieve the dissociation of *in vitro* SSC progenies in the xenovascular model, xenovascular lumens were gently washed with PBS and digested with 1 mg/mL of collagenase/dispase (Sigma-Aldrich) for 10 min at 37 °C. After enzymatic digestion, cells were resuspended in PBS and gently mixed via the pipetting action 10 times, followed by centrifugation at 300 × *g* for 5 min. Harvested cells were resuspended in PBS containing 10% FBS, and probed with anti-mouse PDPN-APC conjugate and anti-human CD31-FITC conjugate. Cell sorting was performed using FACS Aria II (BD Bioscience, San Jose, CA, USA).

### HUVEC proliferation assay

HUVECs were seeded onto a 96-well plate (6.25 × 10^3^/cm^2^) and cultured in complete EGM-2 medium. After pre-culture for 24 h, the culture medium was replaced with the complete EGM-2, non-conditioned medium, and SSC progeny conditioned medium. Cell proliferation was assessed using the WST-8 assay based Cell Counting Kit-8 (DOJINDO LABORATORIES, Kumamoto, Japan).

### HUVEC scratch assay

HUVECs were seeded onto a 24-well plate (0.52 × 10^4^/cm^2^) and cultured with the complete EGM-2 medium until complete confluency was achieved. HUVEC monolayers were starved of EGM-2 for 3 h and scratched using a sterilized 1 mL micropipette tip. The scratched HUVEC monolayers were cultured using a non-conditioned medium or SSC progeny conditioned medium for 24 h. At 0 h and 24 h, microscopy-based images were obtained using inverted optical microscopy (CKX53, Olympus), and the covered area was analyzed using Image J 1.46r.

### Protein array

Angiogenic regulators profiled in the conditioned medium were analyzed using the Proteome Profiler Mouse Angiogenesis Array Kit (ARY015; R&D Systems, Minneapolis, MN, USA). We loaded the conditioned medium (1 mL) onto the array membrane, as per the manufacturer’s instructions. We used ECL Prime (GE Healthcare, Little Chalfont, United Kingdom) as a horseradish peroxidase substrate, and detected chemiluminescence signals using the Light Capture II system (Atto Corporation, Tokyo, Japan). Quantification analysis was performed using Image J 1.46r.

### Statistical analysis

Quantitative data are depicted as mean ± standard deviation values of the mean (SDM) or mean ± standard error values of the mean (SEM). Representative data from at least 3 independent experiments are shown for immunohistochemistry (IHC) and ICC images. Two-group comparisons were made using the unpaired Student’s *t*-test. Multi-group comparisons were made using one-way ANOVA and Tukey’ multiple comparison test or two-way ANOVA and Sidak’s multiple comparison test. Statistical analyses were performed using GraphPad Prism 5 (GraphPad Software, San Diego, CA, USA).

## Acknowledgements

This study was supported by grants-in-aid provided by the Japanese Ministry of Education, Culture, Sports, Science, and Technology (Grant No. 17H05073: S. T and Grant No. 19K08853: A. K), the National Center for Geriatrics and Gerontology (NCGG, the Research Funding for Longevity Sciences, Grant No. 30-11: A. K), the Takeda Science Foundation (S. T), and the SENSHIN Medical Research Foundation (S. T).

## Author contributions

S.T. designed and performed the research, analyzed the data, and wrote the manuscript. M.M., Y.K., W.F., and K.O. performed the experiments. N.S., S.O., A.S., and T. K. analyzed the data. A.K., A.T., K.S-I., and T.M. contributed to the study design and supervised the study. T. Kojima and F.H. conceived and supervised the study. All the authors have discussed the results and commented on the manuscript.

## Competing interests

F.H. received research funding from Daiichi Sankyo, Chugai Pharmaceutical Co, Ltd., Astellas Pharma Inc., and MSD. All the other authors state that they have no competing interests to declare.

## Data availability

All data supporting the findings of this study are available from the corresponding authors upon reasonable request.

## Notes

### Summary of Updates

Supplemental files updated.

