## Supplementary_materials for "Periosteum-derived podoplanin-expressing stromal cells regulate nascent vascularization during epiphyseal marrow development"

1. Supplementary figures (Figure. S1-S5)
2. Supplementary tables (Table. S1 and S2)

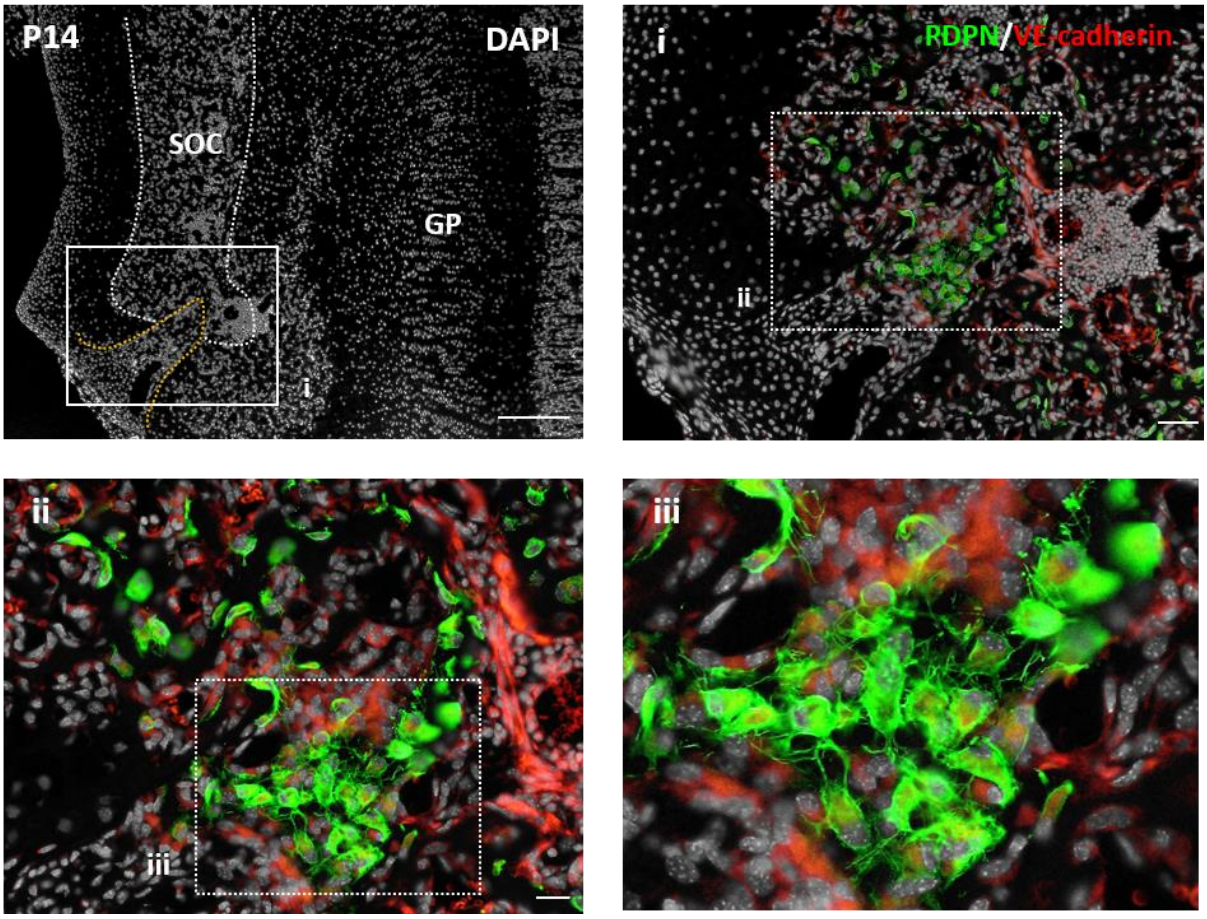


**Figure. S1**
**Proliferative expansion of podoplanin-expressing stromal cells within the penetrating of the periosteal tip.** Representative IHC images of the mouse epiphysis at P14. Cryo-sections of the epiphysis were stained with PDPN, vascular endothelial (VE)-cadherin, and DAPI. Orange dotted-lines indicate the periosteal penetrating tip. Scale bars in the upper left indicate 200 μm. Scale bars in upper right and lower left indicate 50 μm. IHC, immunohistochemistry; P14, postnatal day 14; PDPN, podoplanin; DAPI, 4′,6-diamidino-2-phenylindole; GP, growth plate; SOC, secondary ossification center

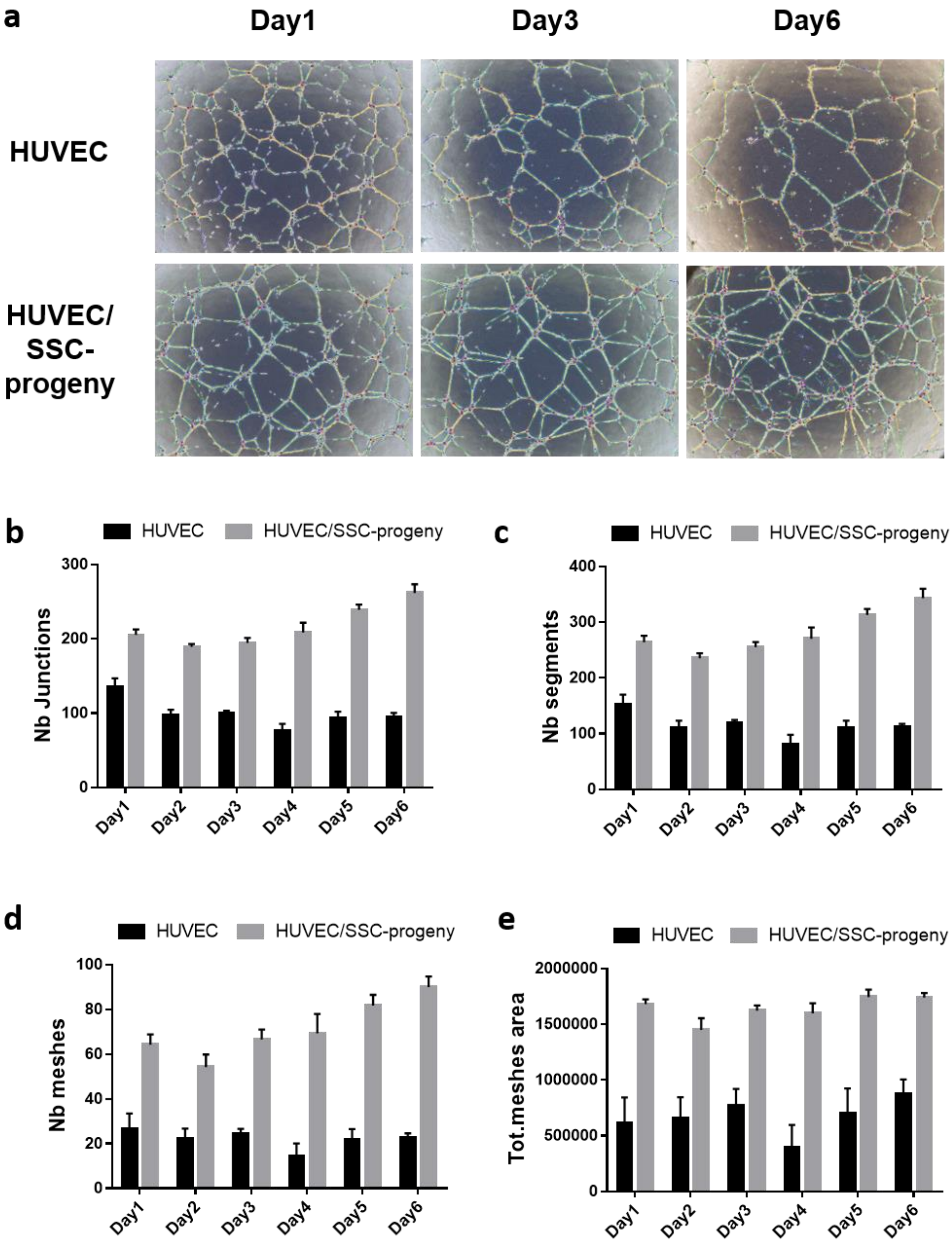


**Figure. S2**

**Long-term study of the xenovascular model co-cultured with HUVECs and PDPN-expressing SSC progenies *in vitro*.** (A) Time series images of HUVEC vascular-like lumens (upper panels) and the xenovascular model co-cultured with HUVECs and PDPN-expressing SSC progenies *in vitro* (lower panels). The day and time above each image indicate the time point from the start of the culture process. (B-E) Quantitative analysis of the integrity of the vascular lumen in the xenovascular model. The parameters for evaluating lumen vascularization, including the number of junctions (B), the number of segments (C), the number of meshes (D), and the total mesh area (E), were measured using an Angiogenesis Analyzer tool. The error bars represent SEMs (n = 5 per group).

| No. | Analyte | Alternative nomenclature | Non-CM (MPI) | SSC-progeny CM (MPI) |
|---|---|---|---|---|
| 1 | ADAMTS1 | METH1 | - | 121.36 |
| 2 | Amphiregulin | AR | - | 96.95 |
| 3 | Angiogenin | ANG | 135.44 | 370.97 |
| 4 | Angiopoietin-1 | Ang-1 | 759.11 | 956.23 |
| 5 | Angiopoietin-3 | Ang-3 | 94.78 | 317.44 |
| 6 | Coagulation Factor III | Tissue Factor, TF | - | 255.02 |
| 7 | CXCL16 | | - | 280.26 |
| 8 | Cyr61 | CCN1, IGFBP-10 | - | 1048.28 |
| 9 | DLL4 | | - | 69.78 |
| 10 | DPPIV | CD26 | - | - |
| 11 | EGF | | - | - |
| 12 | Endoglin | CD105 | - | - |
| 13 | Endostatin/Collagen XVIII | | - | 135.78 |
| 14 | Endothelin-1 | ET-1 | - | - |
| 15 | FGF acidic | FGF-1, ECGF, HBGF-1 | 176.19 | 115.19 |
| 16 | FGF basic | FGF-2 | 244.02 | - |
| 17 | KGF | FGF-7 | - | 100.36 |
| 18 | Fractalkine | CX3CL1 | - | 406.51 |
| 19 | GM-CSF | | - | - |
| 20 | HB-EGF | | - | - |
| 21 | HGF | Hepatopoietin A | - | 304.56 |
| 22 | IGFBP-1 | | - | - |
| 23 | IGFBP-2 | | - | 5355.55 |
| 24 | IGFBP-3 | | - | 1168.82 |
| 25 | IL-1α | IL-1F1 | - | - |
| 26 | IL-1β | IL-1F2 | - | - |
| 27 | IL-10 | CSIF | - | 100.95 |
| 28 | IP-10 | CXCL10, CRG-2 | - | 192.02 |
| 29 | KC | CXCL1, CINC-1, GROα | - | 868.75 |
| 30 | Leptin | OB | - | - |
| 31 | MCP-1 | CCL2/JE | - | 6191.13 |
| 32 | MIP-1α | CCL3 | - | - |
| 33 | MMP-3 (pro and mature form) | | - | 7197.13 |
| 34 | MMP-8 (pro form) | | - | 161.44 |
| 35 | MMP-9 (pro and active form) | | - | - |
| 36 | NOV | CCN3, IGFBP-9 | - | 1706.53 |
| 37 | Osteopontin | OPN | 159.78 | 3614.48 |
| 38 | PD-ECGF | | - | 116.36 |
| 39 | PDGF-AA | | - | 104.36 |
| 40 | PDGF-AB/PDGF-BB | | - | - |
| 41 | Pentraxin-3 | PTX3, TSG-14 | 140.36 | 1826.23 |
| 42 | Platelet Factor 4 | CXCL4, PF4 | - | 176.19 |
| 43 | PIGF-2 | | - | 325.68 |
| 44 | Prolactin | PRL | - | 140.19 |
| 45 | Proliferin | | - | - |
| 46 | SDF-1 | CXCL12 | - | 6855.55 |
| 47 | Serpin E1 | PAI-1 | - | 2194.94 |
| 48 | Serpin F1 | PEDF | - | 281.44 |
| 49 | Thrombospondin-2 | TSP-2 | - | 3969.36 |
| 50 | TIMP-1 | | - | 471.51 |
| 51 | TIMP-4 | | - | - |
| 52 | VEGF | VPF | - | 1208.11 |
| 53 | VEGF-B | VRF | - | 160.78 |

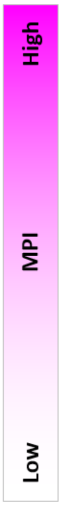


**Figure. S3**

**Quantification of spot intensities in the Proteome Profiler Mouse Angiogenesis Array Kit.** Spot intensity values of the 53 angiogenic regulators present in the Non-CM medium or SSC-progeny CM medium. The MPI range is indicated using the heat distribution depited using the color gradient. Non-CM, EGM2-basal medium supplemented with non-conditioned medium; SSC-progeny CM medium, SSC-progeny conditioned medium; MPI, mean pixel intensity

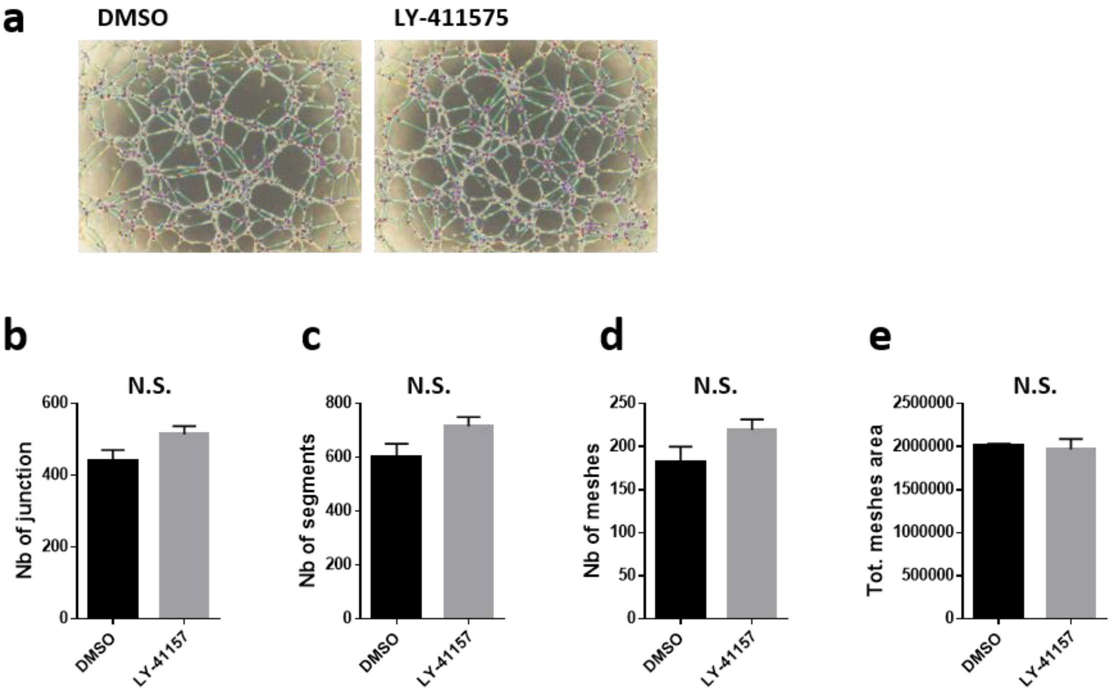


**Figure. S4**

**LY-411575 pretreatment did not affect the morphology and vascular integrity parameters of the xenovascular model.** (A) Representative optical microscopic images of the xenovascular model with vehicle control (DMSO) and LY-411575. (B-E) Quantitative analysis of the vascular integrity. The parameters for evaluating lumen vascularization, including the number of junctions (B), the number of segments (C), the number of meshes (D), and the total mesh area (E), were measured using an Angiogenesis Analyzer tool. N.S. indicates non-significant differences. Statistical analysis was performed using the Student's *t*-test (n = 5 per group). The error bars represent SEMs.

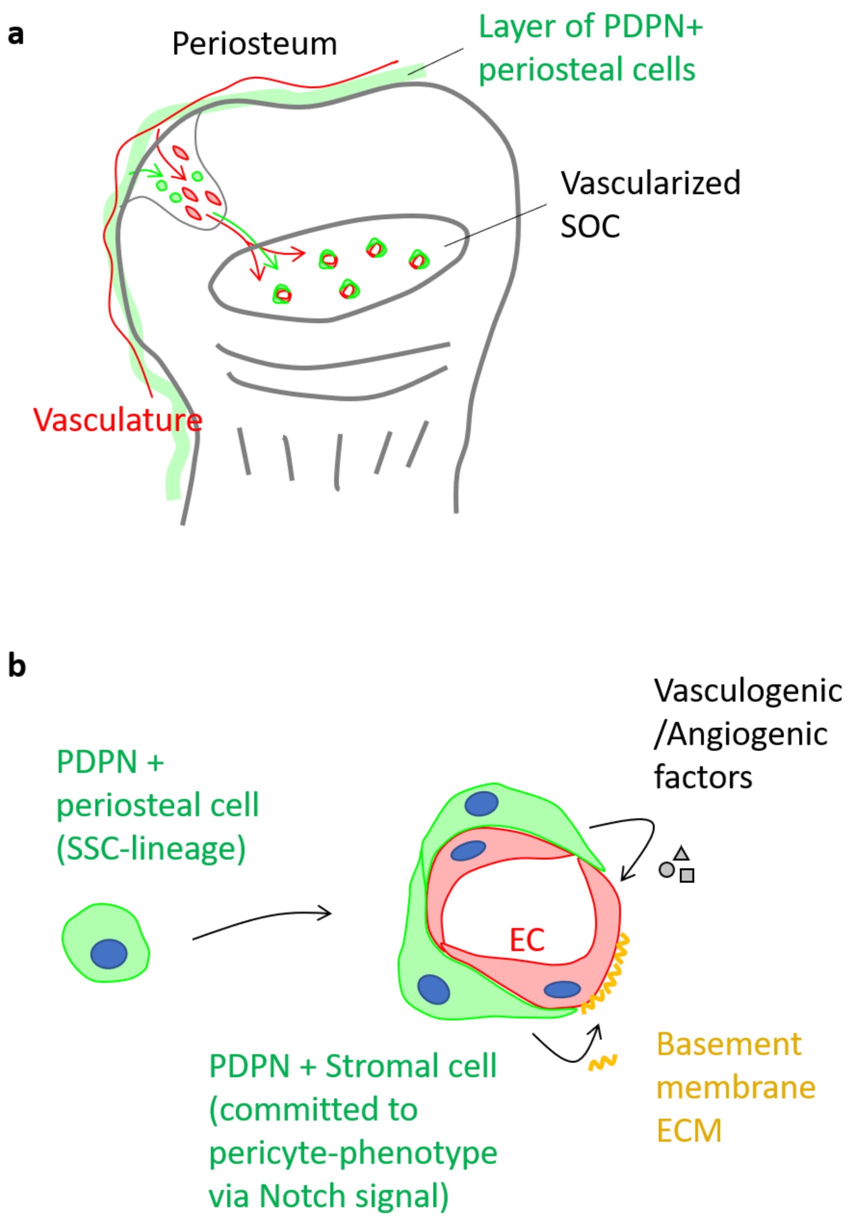


**Figure. S5**

**Graphical model depicting the proposed role of PDPN-expressing stromal cells in epiphyseal marrow development and homeostasis.** (A) Marrow PDPN-expressing stromal cells originate from periosteal cellular components. PDPN-positive periosteal cells invade into the avascular cartilaginous anlage of the postnatal epiphysis and populate the SOC as PDPN-expressing stromal cells. Marrow PDPN-expressing stromal cells behave in a manner similar to the pericytes of the primitive SOC vasculature. (B) Marrow PDPN-expressing stromal cells are the progeny cells of the SSC lineage. Based on the results obtained using the xenovascular model in *in vitro* experiments, we propose that marrow PDPN-expressing stromal cells maintain their vascular integrity by secreting angiogenic factors and vascular basement membrane ECMs. In response to the Notch-mediated interaction with endothelial cells, marrow PDPN-expressing stromal cells commit to the pericyte-phenotype. PDPN, podoplanin; SOC, secondary ossification center; SSC, skeletal stem cell; ECMs, extracellular matrices

**Table S1. Antibodies and experimental conditions used in this study**

| Antibody | Clone | Application/Dilution | Catalog number |
|---|---|---|---|
| Mouse anti-CD4 | Rat mAb (RM4-5) | Lin depletion | 553043, BD Biosciences |
| Mouse anti-CD8 | Rat mAb (53-6.7) | Lin depletion | 553027, BD Biosciences |
| Mouse anti-B220 (Ly5, CD45R) | Rat mAb (RA3-6B2) | Lin depletion | CL8990AP, Cedarlane |
| Mouse anti-TER-119 | Rat mAb (TER-119) | Lin depletion | 553671, BD Biosciences |
| Mouse anti-Ly6G (Gr1) | Rat mAb (RB6-8C5) | Lin depletion | 553123, BD Biosciences |
| Mouse anti-CD11b | Rat mAb (M1/70) | Lin depletion | 553308, BD Biosciences |
| Mouse anti-F4/80 | Rat mAb (Cl.A3-1) | Lin depletion | T-2008, BMA Biomedical |
| Mouse anti-CD71 | Rat mAb (C2) | Lin depletion | 553264, BD Biosciences |
| Mouse anti-PDPN-APC | Hamster mAb (8.1.1) | FCM, PDPN-enrichment | 337022, Biolegend |
| Mouse anti-PDPN | Hamster mAb (8.1.1) | IHC, ICC/1:200 | sc-53533, Santa Cruz Biotechnology |
| Mouse anti-VE-cadherin | Rabbit pAb | IHC, ICC/1:200 | ab32570, Abcam |
| Mouse anti-NG2 | Rabbit pAb | IHC, ICC/1:250 | AB5320, Millipore |
| Mouse anti-αSMA | Mouse mAb (1A4) | IHC, ICC/1:200 | ab7817, Abcam |
| Mouse anti-PDGFRβ | Raabbit mAb (Y92) | IHC, ICC/1:200 | ab32570, Abcam |
| Mouse anti-COL4 (type IV collagen) | Rabbit pAb | ICC/1:200 | ab19804, Abcam |
| Mouse anti-laminins | Rabbit pAb | ICC/1:200 | ab7463, Abcam |
| Human anti-CD31 FITC | Mouse mAb (WM59) | FCM | 11-0319-42, Thermo Fischer Scientific |
| Mouse anti-hematopoietic lineage cocktail-PerCP Cy5.5 | Rat mAb cocktail | FCM | 561317, BD Biosciences |
| Mouse anti-CD31-PerCP Cy5.5 | Rat mAb (390) | FCM | 102419, Biolegend |
| Mouse anti-CD45-PerCP Cy5.5 | Rat mAb (30-F11) | FCM | 550994, BD Biosciences |
| Mouse anti-CD51-PE | Rat mAb (RMV-7) | FCM | 104106, Biolegend |
| Mouse anti-CD150-Alexa488 | Rat mAb (TC15-12F12.2) | FCM | 115916, Biolegend |
| Mouse anti-Sca-1-APC-Cy7 | Rat mAb (D7) | FCM | 108126, Biolegend |
| Mouse anti-CD90.2-BV605 | Rat mAb (53-2.1) | FCM | 140318, Biolegend |
| Mouse anti-CD249-PE-Cy7 | Rat mAb (6C3) | FCM | 108314, Biolegend |
| Mouse anti-CD105-Alexa488 | Rat mAb (MJ7/18) | FCM | 120406, Biolegend |
| Mouse anti-CD200-BV421 | Rat mAb (OX-90) | FCM | 565547, BD Biosciences |
| 7AAD | Viability Staining Solution | FCM | 420404, Biolegend |

### Table S2. qPCR probes used in this study

| Gene name | Assay Name | Ref Seq Number | Label |
|---|---|---|---|
| *Col4a1* | Mm.PT.58.31838522 | NM_009931 | FAM |
| *Col4a2* | Mm.PT.58.28692624 | NM_009932 | FAM |
| *Lama4* | Mm.PT.58.7107286 | NM_010681 | FAM |
| *Lama5* | Mm.PT.58.10095314 | NM_001081171 | FAM |
| *Nid1* | Mm.PT.58.7397156 | NM_010917 | FAM |
| *Nid2* | Mm.PT.58.9823169 | NM_008695 | FAM |
| *Col1a1* | Mm.PT.58.7562513 | NM_007742 | FAM |
| *Col3a1* | Mm.PT.58.13848686 | NM_009930 | FAM |
| *Actb* | Mm.PT.39a.22214843.g | NM_007393 | HEX |
